# Metagenomic survey of antimicrobial resistance (AMR) in Maryland surface waters differentiated by high and low human impact

**DOI:** 10.1101/2023.06.20.545726

**Authors:** Brandon Kocurek, Shawn Behling, Gordon Martin, Padmini Ramachandran, Elizabeth Reed, Christopher Grim, Mark Mammel, Jie Zheng, Alison Franklin, Jay Garland, Daniel A. Tadesse, Manan Sharma, Gregory H. Tyson, Claudine Kabera, Heather Tate, Patrick F. McDermott, Errol Strain, Andrea Ottesen

## Abstract

In alignment with the One Health paradigm, surface waters are being evaluated as a modality to better understand baseline antimicrobial resistance (AMR) across the environment to supplement existing AMR monitoring in pathogens associated with humans, foods, and animals. Here, we use metagenomic and quasimetagenomic sequence data to describe AMR in Maryland surface waters from developed (high human impact) and natural (low human impact) classifications by the National Land Cover Database (NLCD). Critically important β-lactamase genes were observed in twice as many high human impact zones. All data are available under BioProject PRJNA79347. https://www.ncbi.nlm.nih.gov/bioproject/794347

## Announcement

Antimicrobial resistance is recognized as a growing global threat. Surface waters are key integrators between humans, animals, agriculture, and the environment and as such may serve as a valuable modality by which to understand incidence, trends, and drivers of various AMR determinants across the environment. In alignment with One Health strategic planning, the National Antimicrobial Resistance Monitoring System (NARMS) has expanded its traditional whole genome sequencing (WGS) and antimicrobial susceptibility (AST)-based monitoring of AMR of human and animal pathogens to include a pilot evaluation of the important environmental compartment of surface water.

Site selection for surface water collection was determined using land use categories defined by the National Land Cover Database (NLCD 2019) of the United States Geological Survey (USGS). This approach provides a methodology by which to potentially separate “native” AMR in aquatic ecologies from anthropogenically influenced AMR. Twenty liters of water from high (n=15) and low (n=15) impact sites were collected using dead-end ultrafiltration according to previously described methods <u>Dead-end Ultrafiltration Water Collection (protocols.io)</u>. DNA was extracted from replicates of microbiologically enriched and culture-independent samples and sequenced on the Illumina Next Seq 2000 (P3 300 Cycle). Approximately 20 million reads per sample were used with taxonomic and AMR annotation tools (AMR++, AMRFinderPlus, CARD, FDA K-mer[1-5]) to describe bacterial taxa and antimicrobial resistance genes (ARGs) of public health importance to NARMS and veterinary monitoring efforts according to previously described methods[6-9].

Thirty-three ‘critically important’ ARGs from a NARMS watchlist were identified in enriched (quasimetagenomic) samples from 86% (26/30) of sampling sites. These genes confer resistance to: Polymyxin (1), Macrolide (5), β-lactam (15), and Fluoroquinolone (12) classes. From the metagenomic (culture independent) sequence data, only 5 critically important ARGs were detected in 20% (6/30) of sampling sites (Table 1). These results are consistent with previous work that demonstrated that for detection of critically important ARGs from surface water[8] or sediment[10], it is important to include either microbiological enrichment or molecular enrichment. Interestingly, β-lactam resistance genes were prevalent in twice as many of the high impact sites compared to low impact sites (Figure 1).

**Table 1.** Enhanced capacity of quasimetagenomic data to describe important AMR in surface water. From a list of critically important NARMS monitoring targets, the following genes were observed in metagenomic and quasimetagenomic (enriched for 24 H @ 37° in Buffered Peptone Water) data. No critically important ß-lactams were observed in metagenomic data. Genes were annotated using the CARD database. Genes observed at 100% identity across 100% of length are presented here. This table demonstrates the enhanced AMR monitoring capacity that quasimetagenomic data can provide.

| Metagenomic Data | Quasimetagenomic Data |
| --- | --- |
| **Polymyxin**<br>*mcr 3.1* | **Polymyxin**<br>*mcr 3.1* |
| **Macrolide**<br>*mphE, mphF, msrE* | **Macrolide**<br>*acrB, mphA, mphE, mphG, msrE* |
| **Flouroquinolone**<br>*oqxB* | **Flouroquinolone**<br>AAC(6')-Ib-cr, *oqxB*, QnrB1, QnrB19, QnrB2, QnrB38, QnrB6, QnrB7, QnrB9, QnrS1, QnrS2, QnrS4 |
| **Betalactam** | **Betalactam**<br>CMY-2, CMY-30, CMY-4, CMY-43, CTX-M-101, CTX-M-15, CTX-M-3, CTX-M-55, FOX-5, KPC-2, SHV-12, SHV-30, SHV-7, TEM-15, TEM-207 |

**Figure 1.**
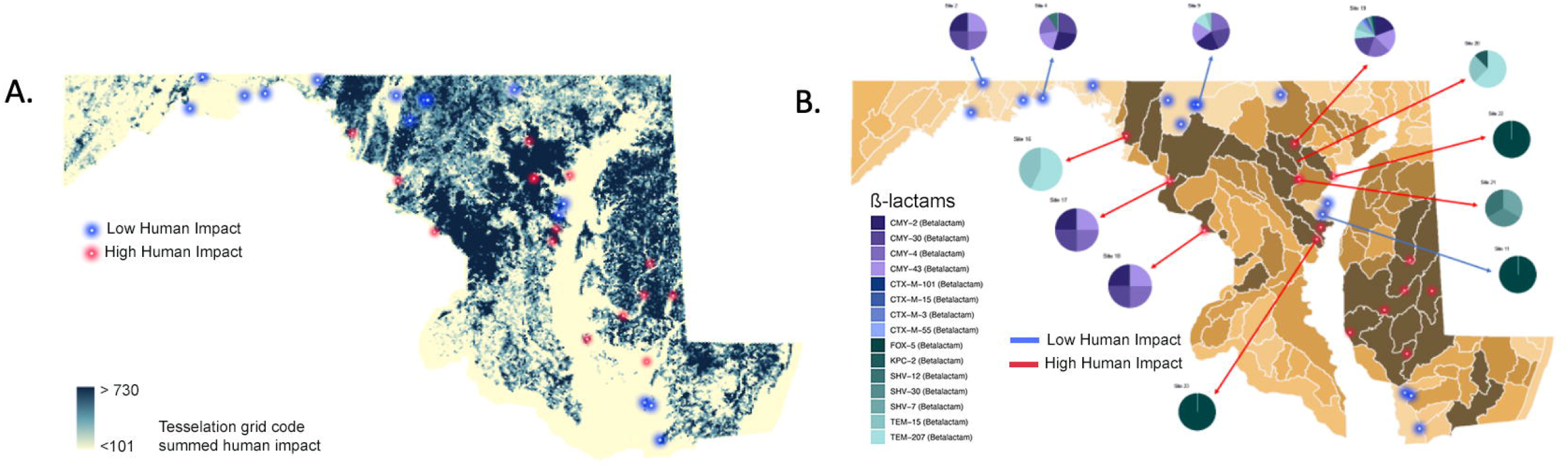
Low and High Human Impact Sites Across Maryland. **Figure 1A**. Land use categories defined by the United States Geological Survey (USGS) were used to select 30 sites across the State of Maryland representing high (red dots) and low (blue dots) “human impact” zones. Our sampling design focused on reclassifying the National Land Cover Database (NLCD 2019; 30 m resolution) into sites representing high intensity of human development contrasted to natural areas. Site selection and all landcover analyses were conducted using ArcGIS Pro 2.9.3 **Figure 1B**. Critically important ß-lactams genes found in quasimetagenomic next-generation sequencing (qmNGS) data annotated using the Comprehensive Antibiotic Resistance Database (CARD) are shown here. Twice as many sites in the high human impact zones were positive for a wide range of ß-lactams compared to the low impact. Interesetingly however – similar ß-lactam resistance determinants were seen across all sites.

Additional foci of the monitoring effort included description of bacterial species most highly correlated with global AMR mortality, NARMS and Veterinary Laboratory Investigation and Response Network (Vet-LIRN) monitoring targets, and species monitored by the European Antimicrobial Resistance Surveillance network of veterinary medicine (EARS-Vet)[11] (Supplementary Table 1). Most of these taxa were not detectable in metagenomic data at the depth of sequencing conducted here and even in enriched (quasimetagenomic) samples, they comprised less than 1% of the data. These data represent some of the first of many that will be contributed by the NARMS surface water monitoring initiative to support evaluation of metagenomic monitoring efforts to better inform agency decisions from a one health perspective. We demonstrate that biological enrichment (quasimetagenomics) improves sensitivity for detecting ARGs of concern to human and animal health and acknowledge that this approach should be bench-marked against molecular enrichment approaches such as PCR panels, bait enrichments, and CRISPR approaches. Data were contributed to the NARMS water metagenome BioProject PRJNA794347 using the Environmental water metadata ontology with additional fields from the One Health Enteric package.

## References

1. Alcock BP, Raphenya AR, Lau TTY, Tsang KK, Bouchard M, Edalatmand A, et al. CARD 2020: antibiotic resistome surveillance with the comprehensive antibiotic resistance database. Nucleic acids research. 2020;48(D1):D517–D25; doi: 10.1093/nar/gkz935.

2. Doster E, Lakin SM, Dean CJ, Wolfe C, Young JG, Boucher C, et al. MEGARes 2.0: a database for classification of antimicrobial drug, biocide and metal resistance determinants in metagenomic sequence data. Nucleic Acids Research. 2019;48(D1):D561–D9; doi: 10.1093/nar/gkz1010.

3. Feldgarden M, Brover V, Gonzalez-Escalona N, Frye JG, Haendiges J, Haft DH, et al. AMRFinderPlus and the Reference Gene Catalog facilitate examination of the genomic links among antimicrobial resistance, stress response, and virulence. Scientific Reports. 2021;11(1):12728; doi: 10.1038/s41598-021-91456-0.

4. Ramachandran P, Mammel M, Ottesen A, Pava-Ripoll M. MitochonTrakr: a reference collection of high-quality mitochondrial genomes for detecting insect species in food products. Mitochondrial DNA Part B. 2019;4(1).

5. Souvorov A, Agarwala R. SAUTE: sequence assembly using target enrichment. BMC Bioinformatics. 2021;22(1):375; doi: 10.1186/s12859-021-04174-9.

6. Cagle R, Ramachandran P, Reed E, Commichaux S, Mammel MK, Lacher DW, et al. Microbiota of the hickey run tributary of the Anacostia river. Microbiol Resour Announc. 2019;8(12):e00123–19.

7. Durigan M, Murphy HR, Silva AJd. Dead-End Ultrafiltration and DNA-Based Methods for Detection of Cyclospora cayetanensis in Agricultural Water. Applied and Environmental Microbiology. 2020;86(23):e01595–20; doi: doi:10.1128/AEM.01595-20.

8. Ottesen A, Kocurek B, Ramachandran P, Reed E, Commichaux S, Engelbach G, et al. Advancing antimicrobial resistance monitoring in surface waters with metagenomic and quasimetagenomic methods. bioRxiv. 2022:2022.04.22.489054; doi: 10.1101/2022.04.22.489054.

9. Smith CM, Hill VR. Dead-End Hollow-Fiber Ultrafiltration for Recovery of Diverse Microbes from Water.Applied and Environmental Microbiology. 2009;75(16):5284–9; doi: doi:10.1128/AEM.00456-09.

10. Gweon HS, Shaw LP, Swann J, De Maio N, AbuOun M, Niehus R, et al. The impact of sequencing depth on the inferred taxonomic composition and AMR gene content of metagenomic samples. Environmental Microbiome. 2019;14(1):7; doi: 10.1186/s40793-019-0347-1.

11. Mader R, Damborg P, Amat J-P, Bengtsson B, Bourély C, Broens EM, et al. Building the European Antimicrobial Resistance Surveillance network in veterinary medicine (EARS-Vet). Eurosurveillance. 2021;26(4):2001359; doi: doi:https://doi.org/10.2807/1560-7917.ES.2021.26.4.2001359.

